# *Bradyrhizobium* and the soybean rhizosphere: Species level bacterial population dynamics in established soybean fields, rhizosphere and nodules

**DOI:** 10.1101/2024.03.15.585214

**Authors:** Sukhvir K. Sarao, Vincent Boothe, Bikram K. Das, Jose L. Gonzalez Hernandez, Volker S. Brözel

## Abstract

*Bradyrhizobium* fixes nitrogen symbiotically with soybean and is an agriculturally significant bacterium. Much is known about the *Bradyrhizobium* species that nodulate soybeans. Conversely, prevalence of *Bradyrhizobium* in soil and the rhizosphere is known only to the genus level as culture independent approaches have provided only partial 16S rRNA gene sequences, so that nodulating and non-nodulating species could not be distinguished. To track which species in bulk soil proliferate in the rhizosphere, and then nodulate, we sought to study population dynamics of *Bradyrhizobium* in soybean fields and rhizosphere at the species level. Recent advances in Oxford Nanopore Technologies provided us with higher fidelity and increased number of reads which enabled us to track *Bradyrhizobium* populations at the species level. We found evidence for 74 species of *Bradyrhizobium* within a community of 10,855 bacterial species in bulk soil and rhizosphere from three different soybean fields in South Dakota. The most predominant species in bulk soil and rhizosphere included *B. liaoningense, B. americanum,* and *B. diversitatus*, however none of these were isolated from nodules. Isolates from nodules included *B. japonicum, B. elkanii* and *B. diazoefficiens.* These nodulators also maintained populations in bulk soil and rhizosphere, although they were not the most prevalent *Bradyrhizobium.* Our findings reveal the rich diversity and community dynamics of *Bradyrhizobium* species in soybean field soil as well as in the rhizosphere. Our results showed that many species of the genus maintain populations in soybean field soil, even in the long-term absence of potential nodulating partners.

## Introduction

Biological nitrogen fixation is a promising solution to replace the use of synthetic fertilizers for cultivation of legumes such as soybean (Howard and Rees 1996). Soybean varieties form symbiotic associations with nitrogen-fixing bacteria of the genus *Bradyrhizobium*. *Bradyrhizobium* are attracted to the soybean root by isoflavones present in soybean root exudates (Subramanian et al 2006). A series of crosstalk events between soybean and *Bradyrhizobium* lead to localized root tissue differentiation that allows entry of the bacterium into an infection thread in the root (Schultze and Kondorosi 1998). This infection thread develops into a nodule that provides nutrients and space to the *Bradyrhizobium* which fixes nitrogen for the plant (Buhian and Bensmihen 2018, Mylona et al 1995). The soybean root employs selection mechanisms that permit only compatible *Bradyrhizobium* to infect the root (Poole et al 2018).

*Bradyrhizobium* which are incompatible with a particular soybean variety and therefore unable to nodulate would be at a disadvantage as they remain outside the roots. Furthermore, *Bradyrhizobium* are best known as nodule occupying or associative nitrogen fixers (Parker 2015). However, a recent meta-study has shown that *Bradyrhizobium* is the predominant genus in soils across the globe (Delgado-Baquerizo et al 2018). Many of these soils contain few or no legumes, so *Bradyrhizobium* must display population fitness outside of nodules. While much is known about nodulation and physiology of *Bradyrhizobium* inside nodules, little is known about the ecophysiology of the genus in soil. *B. japonicum, B. diazoefficiens* and *B. elkanii* are common nodulators of soybean (Nakei et al 2022), but other *Bradyrhizobium* have been isolated. Furthermore, *Ensifer* (Li et al 2011), and *Sinorhizobium* (HA and SALEH-RANSTIN 2002) have been isolated from nodules in sodic soils. To date, 89 *Bradyrhizobium* species have been reported in the List of Prokaryotic Names with Standing in Nomenclature (https://lpsn.dsmz.de last accessed 24 February 2024). A genomotaxonomic study predicted the genus to comprise approximately 800 species (Ormeño-Orrillo and Martinez-Romero 2019). This indicates that the genus is largely uncharacterized and that much remains to be discovered. Due to the limitations of high-throughput DNA sequencing, bacterial diversity studies rely on partial sequences of the 16SrRNA gene pool, e.g. the V3 to V4 region, supporting allocation of reads to genus at best. To date, evidence for *Bradyrhizobium* in soil has been based on partial sequences which only allow allocation to genus, so that no information is available on distribution of specific species in the environment. The prevalence of specific *Bradyrhizobium* species that can or cannot nodulate soybeans, cannot be deduced from partial sequence-based data, causing the putative presence of nodulators to be masked by lack of information. To obtain information on prevalence of specific species will require more information per sequence variant, for example, complete sequencing of 16SrRNA genes. Full 16SrRNA gene data would be instrumental in revealing which *Bradyrhizobium* in bulk soil seed the rhizosphere, and which of these nodulate soybean. This would show which of the *Bradyrhizobium* species are nodulating, and which are non-nodulating under field conditions with high accuracy.

In the soybean rhizosphere, nodulating *Bradyrhizobium* must compete with other strains and other genera present before forming nodules. Following seed germination, roots grow and extend into bulk soil, providing an expanding rhizosphere (Sun et al 2021). Bulk soil bacteria able to benefit from root exudates will seed the rhizosphere community, while other bulk soil bacteria will not. Thus, growing roots cause shifts in community composition (Sugiyama 2019) (Hassan et al 2019), opening the door to interactions among different species. So, *Bradyrhizobium* would encounter competing species. To better understand competition among bacteria in the rhizosphere, it is imperative to characterize rhizosphere communities at the species level. While reductionist interaction experiments have been performed using two strains (Herschend et al 2022) (Galet et al 2015) (Du et al 2024),(Lyng et al 2024) species level data of rhizosphere in field conditions is lacking. Moreover, species level characterization would improve understanding of how *Bradyrhizobium* inoculants perform in the rhizosphere community and how this impacts nodulating capacity.

In the competitive environment of the rhizosphere, nodule forming *Bradyrhizobium* must compete with numerous other bacteria seeded from bulk soil. We asked: Which *Bradyrhizobium* species occur in soybean field bulk soil, which of these occur in the soybean rhizosphere, and finally which of those end up in nodules? We hypothesized that the nodulating species *B. japonicum, B. diazoefficiens* and *B. elkanii* would predominate in soils under long term soybean cultivation. To characterize microbial communities to species level, we obtained full 16SrRNA gene sequences from bulk soil and soybean rhizosphere at three different field sites in South Dakota, using Oxford Nanopore Technologies (ONT) long read sequencing on MinION flowcell. The results did not support our hypothesis. Surprisingly, we found a diversity of *Bradyrhizobium* species, as well as many members of other associative nitrogen-fixing genera occurring in the bulk soil, to be enriched in the rhizosphere of soybeans grown in the different soil types. As expected, nodules formed in this diverse diazotroph environment contained only *B. japonicum, B. diazoefficiens* or *B. elkanii*.

## Materials and Methods

### Sample collection and processing

Samples were taken at three of eight locations as part of a larger isolation study in South Dakota on June 12, 2023. Sample 5 was at coordinates 41.14667 N, 96.788889 W (Sinai Kranzburg Barnes), sample 6 at 44.108889 N, 96.768333 W (Sinai Kranzburg Barnes), and sample 8 at 44.306944 N, 96.671111 W (Alluvial soils undifferentiated). Soybean plants (5 per location) were dug up and placed into sterile plastic bags. Bulk soil samples taken next to the plants were placed into separate plastic bags. After transportation to the laboratory on ice, soil samples were frozen at −80°C for later DNA extraction. Shoots were cut off plants and roots were washed in sterile water to remove the adhering soil. Nodules (3 per plant) were harvested using sterile tweezers and stored at – 80°C until further use. The remaining nodules were removed to avoid inclusion of nodule bacteria in the rhizosphere preparation. The rhizosphere material was then obtained from these roots by applying multiple wash steps and sonication as described previously (White et al 2015). The first sonication extract was discarded. Fractions from the second and third sonication were pooled and frozen at −80°C for later DNA extraction. DNA was extracted from bulk soil samples using the DNeasy Power Soil Pro Kit (47014 QIAGEN, Hilden, Germany). DNA from the rhizosphere was extracted using the DNeasy UltraClean Microbial Kit (12224-50 QIAGEN, Hilden, Germany). DNA extracts were stored at −80°C. Bulk soil samples were submitted for chemical analysis to Midwest Laboratories (Omaha, NE).

### Isolating *Bradyrhizobium* from the nodules

To obtain isolates from nodules (15 per location), nodules were individually thawed by immersion in 500 µl R2 broth in 1.5 ml microfuge tubes overnight. Nodules were placed onto paper towels soaked in 70% ethanol and rubbed to remove residual material. Nodules were then rinsed three times in 500 µl sterile water. Nodules were transferred into 1.5ml microfuge tubes and crushed using sterile tweezers or a toothpick. The nodule extract was resuspended in 500 µl sterile water, a loop-full streaked onto BJSM selective media (Tong and Sadowsky 1994), and incubated at 28°C for 4-6 days until colonies became visible. To exclude rapidly growing bacteria (non-*Bradyrhizobium*), plates were inspected after 2 days, and visible colonies labeled for later exclusion. Colonies appearing by day four or later were re-streaked onto R2A plates containing cycloheximide (100 mg/L to suppress fungal growth) to obtain pure cultures. Isolates were selected based on time to emergence and gram-negative small rods were selected for confirmation by 16SrRNA gene sequencing.

### DNA extraction and phylogeny of nodule isolates

DNA was extracted from nodule isolates using DNeasy UltraClean Microbial Kit (12224-50 QIAGEN, Hilden, Germany. The 16S rRNA gene was PCR amplified using primers 27F (AGAGTTTGATYMTGGCTAG) and 1492R (TACGGYTACCTTGTTACGACTT) and submitted for Sanger dideoxy chain termination sequencing. Partial sequence results were submitted to BLAST search (Altschul et al 1990) to tentatively allocate isolates to genus. The 16SrRNA genes of putative *Bradyrhizobium* isolates were reamplified, quantified using the Qubit^TM^ dsDNA BR Assay Kit (Q32850) and a Qubit spectrofluorometer (Qubit 3.0, Thermo Fisher Scientific, USA), and submitted for complete sequencing (Plasmidsaurus Inc., Eugene, OR, USA). Relatedness of isolate sequences to genome-derived sequences from NCBI was determined by alignment using MUSCLE, and a tree was generated using the maximum likelihood method in MEGA version 11 (Tamura et al 2021).

### Sequencing and analysis of full length 16S rRNA gene data

The library preparation and sequencing were done in the SDSU Genomics Sequencing Facility. Sequenicng libraries were prepared using (ONT) −16S Barcoding Kit (SQK-16S024) following the manufacturer’s protocol (ONT, Oxford, UK) using 10 ng of DNA from each sample. Primer sequences were: Forward 16S primer: 5’ ATCGCCTACCGTGAC - barcode - AGAGTTTGATCMTGGCTCAG 3’; Reverse 16S primer: 5’ ATCGCCTACCGTGAC - barcode - CGGTTACCTTGTTACGACTT 3’. The sequencing of the libraries was done on a MinION sequencer (ONT, Oxford, UK) using a R9.4.1 flowcell. The demultiplexed sequence data was analyzed using EPI2ME v5.1.9 (ONT 2023). The “.fastq” files were first analyzed using the workflow: “wf-16S”. The workflow was set to use minimap2 for classification. Under the reference option, the ‘ncbi_16s_18s’ was selected as a reference for classifying reads. Furthermore, under the “Taxonomic rank” settings, the Kraken2 (Wood et al 2019) pipeline was set to the level that Bracken estimated the abundance at S (Species) level, changing it from the default setting that uses G (Genus) level estimation. To filter to the reads matching with *Bradyrhizobium*, under the Minimap2 options, TaxID: 374 (Schoch et al 2020) was entered. The remaining setup for the analysis was set to the default. After the analysis was completed, the OTU table and the *Bradyrhizobium* matching read stats were analyzed on R v4.3.2 (Team 2021b) and RStudio (Team 2021a). MicrobiomeAnalyst 2.0 (Lu et al 2023) was used to obtain alpha diversity, beta diversity, percentage abundance, heat tree, and correlations at order and family level. Abundance of nitrogen-fixing genera was extracted from OTU tables using data reported recently (Koirala and Brözel 2021) using Excel. Sequences were deposited at NCBI under BioProject accession number PRJNA1085322.

## Results

ONT’s MinION yielded long reads of 16SrRNA gene, with which we were able to allocate sequences to species level. We found evidence for 74 *Bradyrhizobium* species occurring in the three soil types and enriched in the soybean rhizospheres. Surprisingly, we also found many species of other associative nitrogen-fixing genera, including *Rhizobium, Sinorhizobium,* and *Devosia,* enriched in the rhizosphere. Yet only *B. japonicum, B. diazoefficiens,* and *B. elkanii* were found in nodules. Soils at the three sites were of different types, and had chemically different properties including pH, salinity, and ionic composition (Table 1). Sites 5 and 6 were slightly alkaline while site 8 was acidic.

**Table 1:** Organic matter, ionic composition, pH and salinity of soils from three sampling sites.

| Sample | Organic matter | Phosphorus |  | Potassium | Magnesium | Calcium | Sodium | pH | Cation exchange capacity | % Base saturation (computed) |  |  |  |  | Soluble salts |
| --- | --- | --- | --- | --- | --- | --- | --- | --- | --- | --- | --- | --- | --- | --- | --- |
|  | L.O.I. | P1<br>(Weak Bray)<br>1:7 | P2<br>(Strong Bray)<br>1:7 |  |  |  |  |  |  | %K | %Mg | %Ca | %H | %Na |  |
|  | % | ppm | ppm | ppm | ppm | ppm | ppm | 1:1 | meq/100g | 3.3 | 21 | 75.5 | 0 | 0.2 | mmhos/cm |
| 5* | 5.2 | 65 | 135 | 322 | 629 | 3780 | 9 | 7.6 | 25 | 1.3 | 18 | 80.4 | 0 | 0.3 | 0.6 |
| 6* | 2.4 | 11 | 47 | 106 | 444 | 3291 | 15 | 8.1 | 20.5 | 1.7 | 19.9 | 51.8 | 26.3 | 0.3 | 0.4 |
| 8* | 3.5 | 24 | 38 | 136 | 485 | 2104 | 14 | 5.5 | 20.3 | 3.3 | 21 | 75.5 | 0 | 0.2 | 0.3 |

\* Bulk soil samples taken adjacent to soybean plants from three of eight locations in South Dakota that form part of a larger study. See materials and methods for location details.

To attain an overall view of the bacterial communities, we determined alpha and beta diversities. Alpha diversity (Shannon index) of the rhizosphere bacterial community was lower than in bulk soil (Fig 1 a,b). This indicates that the root exudates of growing soybeans benefited a subset of bulk soil bacteriome to grow. Chao diversity index was higher in rhizosphere of sites 5 and 8 when compared to the bulk soil, indicating detection of taxa that were below detection limit in the bulk soil. This could be because some rare species benefited from root exudates, growing to levels above the detection threshold, while not detected in bulk soil samples. Alpha diversity by both metrics was very similar across the four reps of bulk soil but varied across reps of rhizosphere samples. This indicated that the bulk soil community responded variably to presence of a growing root. Community composition as measured by beta diversity differed substantially between bulk soil and rhizosphere. Despite the differences in soil composition between sites 5 and 6, the respective communities were very similar (Fig 1 c,d). Community composition in the acidic site 8 was substantially different from sites 5 and 6 for both bulk soil and rhizosphere. This indicated that pH may be a primary driver of community composition, compared to other soil characteristics such as phosphate, organic matter etc. (Table 1). The four reps of each bulk soil were closer to each other than the rhizosphere reps, indicating that the beta diversity of rhizosphere samples varied more among samples per site than in bulk soil. This agreed with diverging alpha diversity of rhizosphere samples. Collectively, the data pointed to a highly diverse bulk soil communities that shifted substantially after exposure to growing roots. While soil pH affected community composition, the other chemical metrics as reported in Table 1 did not.

**Figure 1:**
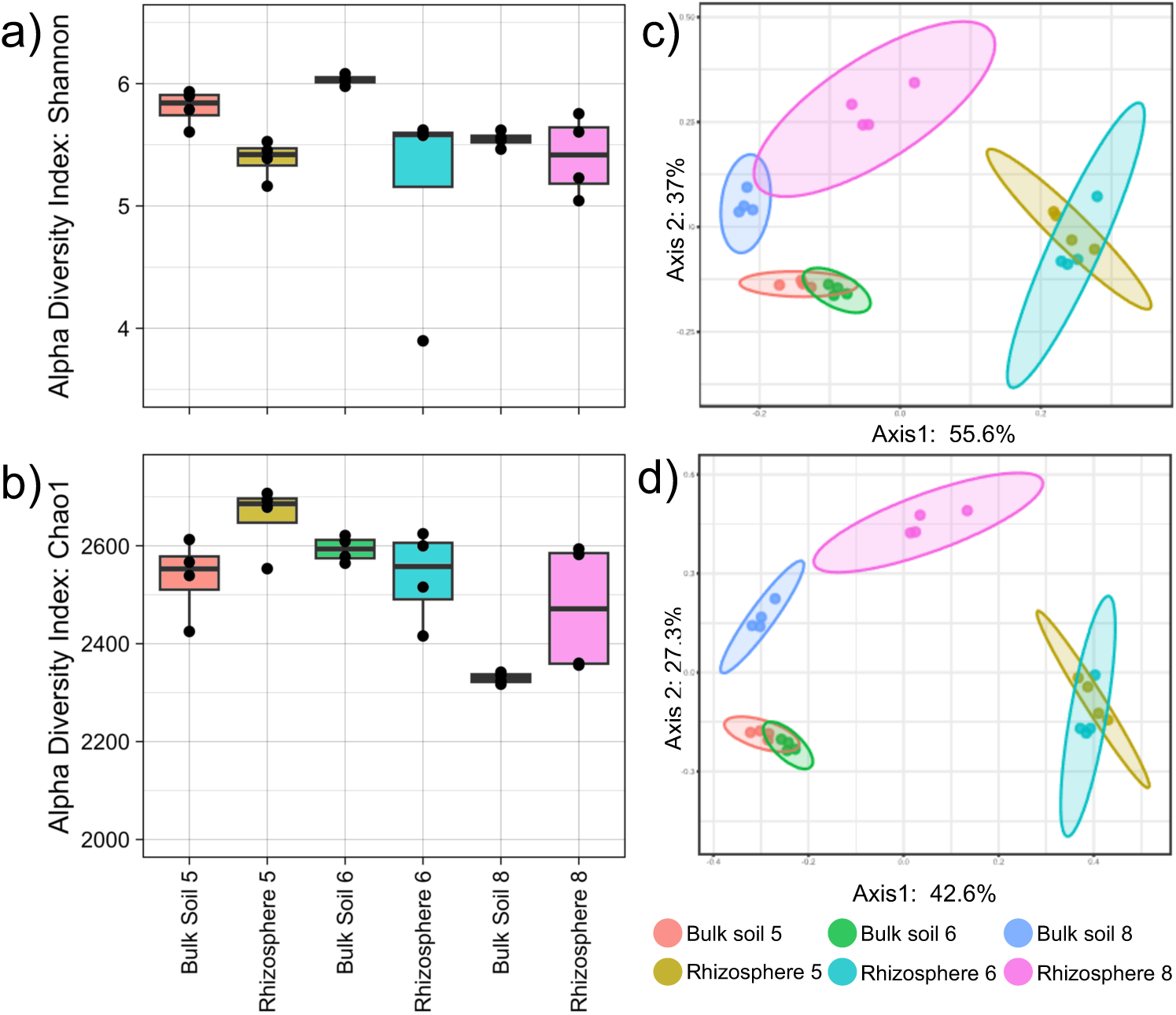
Alpha-diversity derived using the Shannon (a) and Chao1 (b) metrics, and Beta-diversity derived using the Jensen-Shannon divergence (c) and Bray-Curtis index (d) metrics, prepared using Microbiome Analyst.

To visualize shifts in predominance of taxa at the class level, we constructed heat trees (Fig 2a-c). The Phylum Proteobacteria were more abundant in rhizosphere than in bulk soil, while the Firmicutes, Actinobacteria, Chloroflexi, and Acidobacteria were less abundant as indicated by the blue and red lines respectively (Fig 2). The Bacteriodota did not shift much overall. The predominance of several classes shifted substantially in the rhizosphere of all three soils. This was surprising as the three soils comprised different chemical compositions. The classes Alphaproteobacteria, Betaproteobacteria, Gammaproteobacteria, and Nitrospinia were more predominant in rhizosphere, while Deltaproteobacteria, Nitrospira, Clostridia, Bacilli, and Hydrogenophilalia were lower as indicated by the red color of lines (Fig 2). Additional shifts were seen in specific soils, for example Aquificae in sites 5 and 6, and Flavobacteria only in site 5. These results indicate that rhizosphere community composition was influenced more by presence of roots and their exudates, than by soil chemistry.

**Figure 2:**
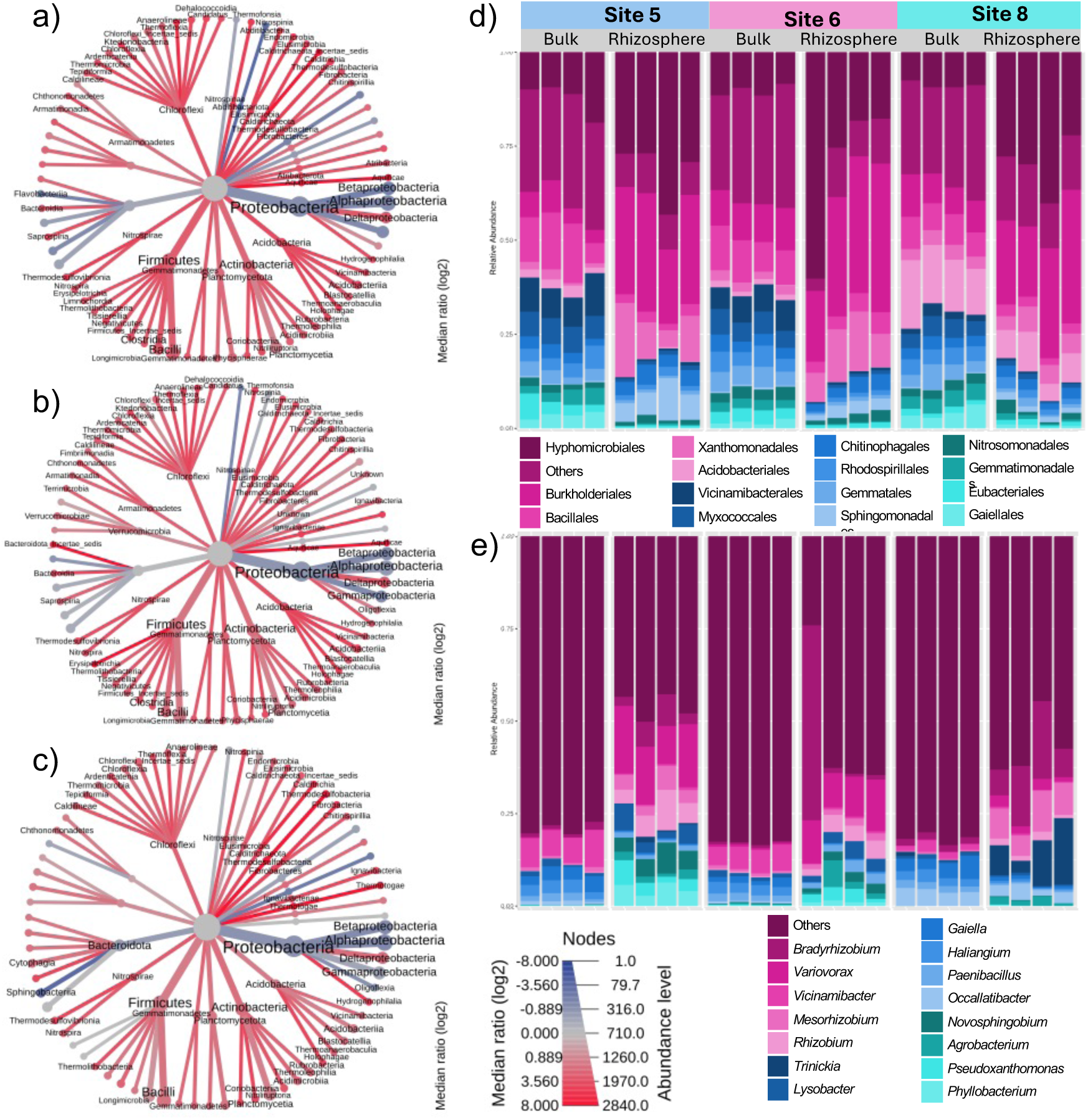
Distribution of taxa in bulk soil versus rhizosphere at the class, order, and genus levels. Relative proportions of classes are shown by heat trees with red lines indicating predominance in bulk soil and blue in rhizosphere (a-c). Proportions of the predominant 15 orders (d) and genera (e) are shown by stacked bar plots. Figures were derived using Microbiome Analyst.

To better understand the rhizosphere-associated increase in proportions of Alpha-, Beta- and Gammaproteobacteria, we visualized the predominant orders and genera (Fig 2 d and e). The most predominant orders in rhizosphere were Hyphomicrobiales (Alphaproteobacteria), Burkholderiales (Betaproteobacteria) and Xanthomonadales (Fig 2d). Rhizosphere Hyphomicrobiales were predominated by *Bradyrhizobium* (1.2% in bulk soil and 10.6% in rhizosphere), *Mesorhizobium* (0.63% in bulk soil and 4.6% in rhizosphere) and *Rhizobium* (0.16% in bulk soil and 4.0% in rhizosphere). One Site 6 rhizosphere sample contained a much higher proportion of *Bradyrhizobium* than the other three reps. Although we attempted to remove all nodules before rhizosphere extraction, some soybean nodules can be partially embedded in the roots, causing nodule tissue containing *Bradyrhizobium* to be left behind in the root.

The predominant rhizosphere member of Burkholderiales was *Variovorax* (0.29% in bulk soil and 9.5% in rhizosphere) and for Xanthomonadales (now Lysobacterales) it was *Lysobacter* (1.07% in bulk soil and 2.58% in rhizosphere). Bacillales, Vicinamibacterales, Acidobacteriales and Myxococcales decreased in the rhizosphere (Fig 2d). *Paenibacillus* decreased from 1.8 to 1.2% while the well-known soil bacterium *Bacillus* decreased from 1.26% in bulk soil to 0.42% in the rhizosphere. *Vicinamibacter* (Vicinamibacterales) decreased from 5.2 to 0.43%, *Gaiella* (Gaiellales) from 3.05 to 0.4% and *Haliangium* (Myxococalles) from 2.7 to 0.37%. As expected, the acidophilic Acidobacteriales (now Terriglobales in the Acidobacteriota) were much more abundant in the bulk soil and rhizosphere in the acidic site 8 compared to the other two sites. *Variovorax* is known to benefit growth of nitrogen-fixing bacteria in the rhizosphere (Flores-Duarte et al 2022). Intriguingly, the predominant rhizosphere genera *Bradyrhizobium*, *Mesorhizobium,* and *Rhizobium* are associative nitrogen fixers of legumes, but soybeans are nodulated only by selected species of *Bradyrhizobium* (Stougaard 2000). These data indicated that associative nitrogen-fixing bacteria and their syntrophic partners, not just those known for crosstalk via isoflavones and nod factors, were enriched in the rhizosphere (Mayhood and Mirza 2021). This prompted us to extract data for associative and free-living nitrogen-fixing genera.

To map community composition of nitrogen-fixing genera in bulk soil versus rhizosphere, we charted the community percentage of nitrogen-fixing bacteria to include associative (nodule-forming), acidophilic associative, and free-living nitrogen fixers, and of *Frankia*. To attain an overall view of nitrogen-fixing bacteria in bulk soil and the rhizosphere, we included all genera identified recently to contain the core nitrogenase genes *nifHDKENB* (Koirala and Brözel 2021). Not all members of these genera are known to contain core nitrogenase genes, so the numbers shown in Fig 3a are likely an overestimation. Yet they point to significant increase of nitrogen fixers in the soybean rhizosphere as seeded by bulk soil bacteriota. In all three sites, *Bradyrhizobium* increased in abundance in rhizosphere, but so did other associative nitrogen-fixing genera such as *Rhizobium, Mesorhizobium, Sinorhizobium, Neorhizobium, Devosia, Ensifer,* and *Shinella* (Fig 3 b-i). The acidophilic associative nitrogen fixers *Trinickia, Burkholderia,* and *Paraburkholderia* were more predominant in the acidic site 8. Yet none of these were obtained from nodules (Fig S1). The other nodulating nitrogen fixers were depleted in acidic soil rhizosphere, with the exception of *Bradyrhizobium* and *Mesorhizobium.* The well-known free living nitrogen fixers *Azospirillum, Azoarcus, Azospira,* and *Paenibacillus* all decreased in rhizosphere, as did *Frankia*. *Herbaspirillum* increased significantly in the acidic soil rhizosphere. Collectively these data show that the soybean roots benefited rhizosphere population growth of diverse associative nitrogen fixers, even if most of these did not occur in nodules. Conversely, free-living nitrogen fixers and *Frankia* were disadvantaged in the rhizosphere.

**Figure 3:**
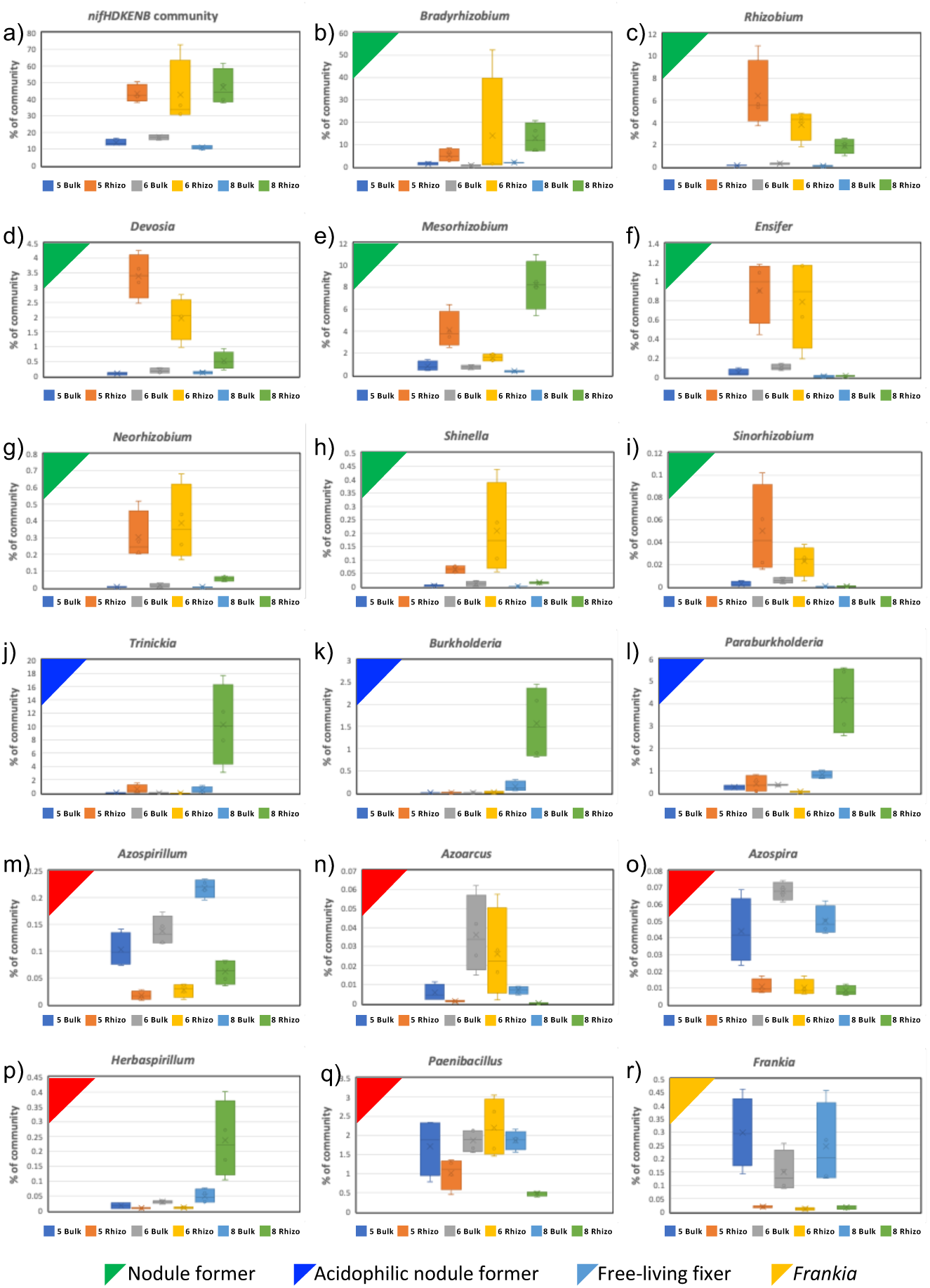
Proportion of entire bacterial community of *nifHDKENB* – bearing genera, associative nitrogen-fixing genera, free-living nitrogen-fixing genera, and *Frankia*.

As associative nitrogen fixer genera all increased in rhizosphere, we sought to identify correlations amongst nitrogen-fixing bacterial families, namely Nitrobacteraceae (includes *Bradyrhizobium*), Rhizobeaceae (includes *Rhizobium, Ensifer, Sinorhizobium,* and *Neorhizobium*), and Devosiaceae (includes *Devosia*). Nitrobacteraceae correlated positively with Rhizobiaceae and Devosiaceae in sites 5 and 8, as indicated by red lines, but no correlation was seen in site 6 at the cut-off of 0.87 (Fig 4). For site 5, Nitrobacteraceae correlated with Rhizobeaceae at 0.943, Nitrobacteraceae with Devosiaceae at 0.92, and for site 8 Nitrobacteraceae with Rhizobeaceae at 0.96, and Nitrobacteraceae with Devosiaceae at 0.89. This indicates that soil parameters other than pH (Table 1) also influence fitness and population dynamics/interactions amongst specific families. The Nitrobacteraceae also correlated positively with multiple other families, including Rhodobiaceae, Caulobacteraceae, and Aurantimonadaceae in site 5, only with Sphingobacteriaceae in site 6, and Calldimonadaceae, Legionellaceae, and Pseudomonadaceae in site 8. Negative correlations were seen with the families Thermomicrobiaceae, Anaerolineaceae, and Geodermatophilaceae in site 5, Desulfosalsimonadaceae, Dehalococcoidaceae, and Calditrichaceae in site 6, and Holophagaceae, Endomicrobiaceae, and Usitatibacteraceae in site 8. This indicated that alkalinity and other soil parameters decreased the correlation capacity of Nitrobacteraceae in soil.

**Figure 4:**
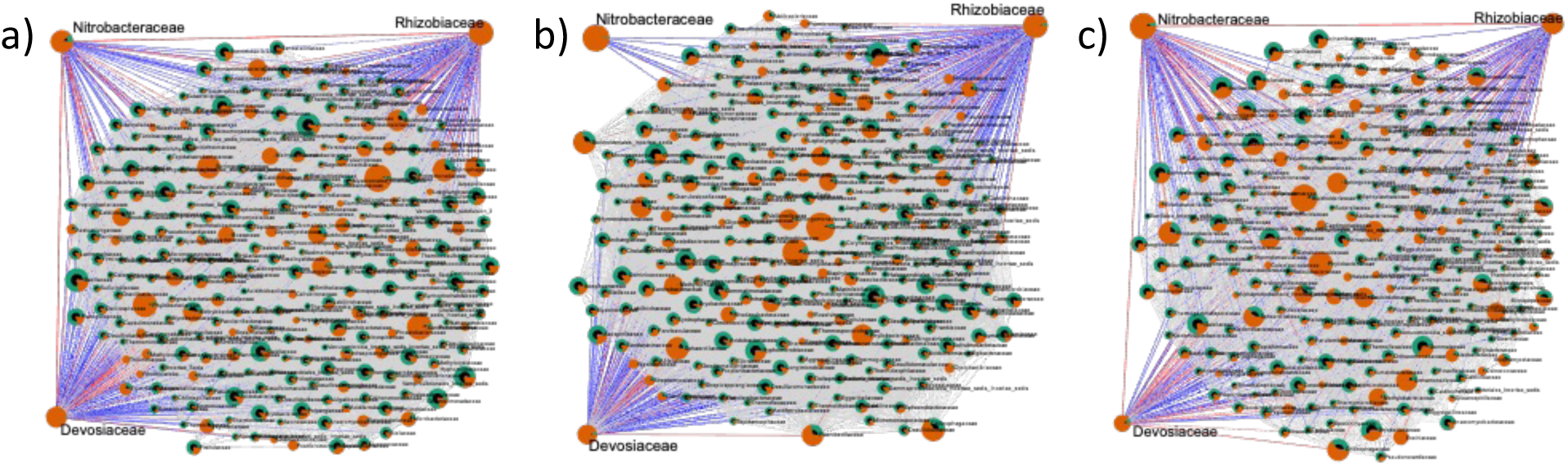
Correlation network derived using SECOM (Pearson 2) at Family level, A) site 5 B) Site6 C) site 8, highlighting Nitrobacteraceae, Rhizobiaceae and Devosiaceae. Red lines show positive correlation > 0.87 at p < 0.005, while blue lines show negative correlations. Green segments represent proportion in bulk soil, and orange in rhizosphere. Figures were derived using Microbiome Analyst.

To determine whether the Hyphomicrobiales correlated with the acidophilic Burkholderiales (includes *Burkholderia*, *Paraburkholderia* and *Trinickia*) in the acidic site 8 versus sites 5 and 6, we derived correlation networks. The Hyphomicrobiales did correlate positively with Burkholderiales in site 8 at 0.95, and at 0.9 and 0.89 in sites 5 and 6 respectively (Fig 5). This positive correlation indicated that both Hyphomicrobiales and Burkholderiales were similarly competitive in the rhizosphere environments in neutral and acidic soils. However, Burkholderiales correlated positively with more orders in acidic soil than in sites 5 and 6. The orders correlating positively with Burkholderiales were Caulobacterales, Sphingomonadales, and Xanthomonadales in all three sites and the ones correlating negatively included Tepidiformales, Thermoanaerobaculales, and Nitrospinales.

**Figure 5:**
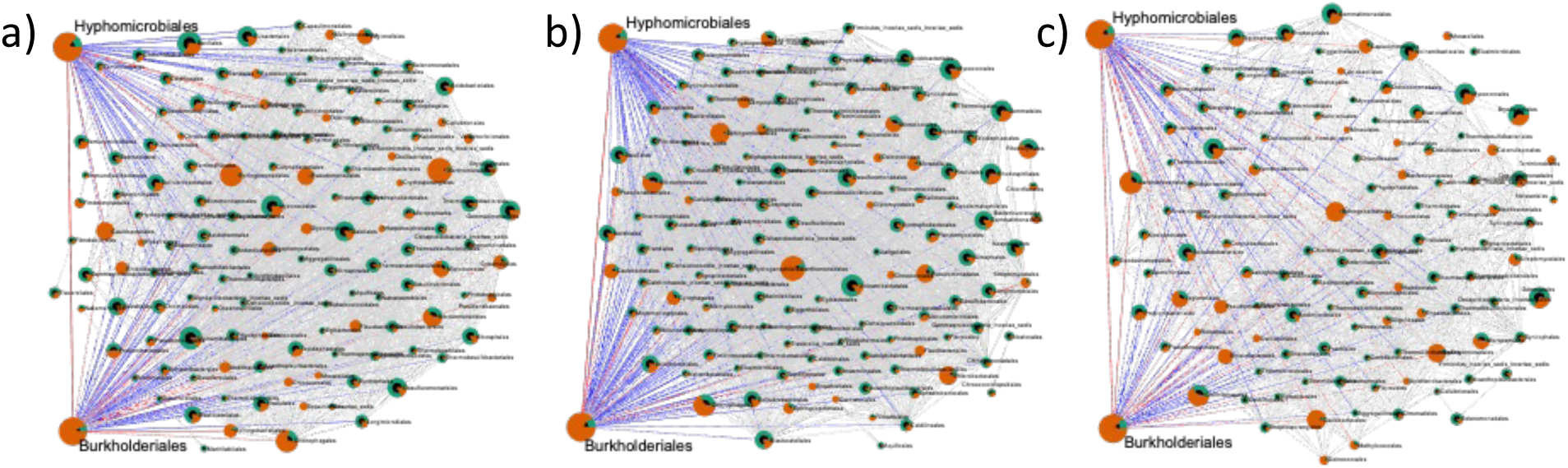
Correlation network derived using SECOM (Pearson 2) at Order level, A) site 5 B) Site6 C) site 8, highlighting Hyphomicrobiales and Burkholderiales. Red lines show positive correlation > 0.9 at p < 0.005, while blue lines show negative correlations. Green segments represent proportion in bulk soil, and orange in rhizosphere. Figures were derived using Microbiome Analyst.

To determine which species of *Bradyrhizobium* occurred in bulk soil, rhizosphere and nodules, we allocated the Nanopore reads to species level and compared these to the full 16S rRNA sequences of nodule isolates. We found evidence of 74 *Bradyrhizobium* species in both bulk soil and rhizosphere, almost all of which occurred at all three sites (Fig 6). In contrast, we only obtained *B. japonicum, B. elkanii,* and one *B. diazoefficiens* isolate from multiple nodules of the multiple plants per site (Fig S1). The acidic site had the highest number of *Bradyrhizobium* species (Fig 6c). In almost all cases, *Bradyrhizobium* were more abundant in rhizosphere than bulk soil, in agreement with genus level data (Fig 3b). *B. lianongense, B. americanum and B. diversitatus* were most predominant as is highlighted by the log scale of Figure 6. In contrast, the three nodulating species comprised 10%, 5%, and 1% of the overall rhizosphere *Bradyrhizobium* community, highlighted by black boxes (Fig 6). These results indicate that the soybean root supported growth of all *Bradyrhizobium* occurring in bulk soil, and not only growth of nodule forming strains (Fig 6). *B. elkanii* was less abundant in site 6 bulk soil, and only one of 5 nodule isolates corresponded to this species. We set out to obtain isolates from 3 nodules per plant to a total of 45 nodules, but only obtained 10 isolates from site 5, 6 from site 6 and 12 from site 8. This was primarily due to overgrowth by fast-growing bacteria. Initial attempts to surface sterilize the field nodules led to apparent killing of the contents of these small nodules. We therefore performed a gentler surface cleaning than is widely prescribed for greenhouse grown soybean nodules. Some species, like *B. ingae* and *B. cenepique,* occurred only in site 8 bulk soil, but were detectable in three rhizospheres, indicating their acidophilic nature. Conversely, species like *B. macuxiense* and *B. denitrificans* were not detected in site 6 rhizosphere, although they were present in bulk soil. Collectively, the data indicate that the soybean root environment supports growth of most *Bradyrhizobium* species occurring in bulk soil, as soybean nodulating species did not constitute the majority of *Bradyrhizobium* in the rhizosphere.

**Figure 6:**
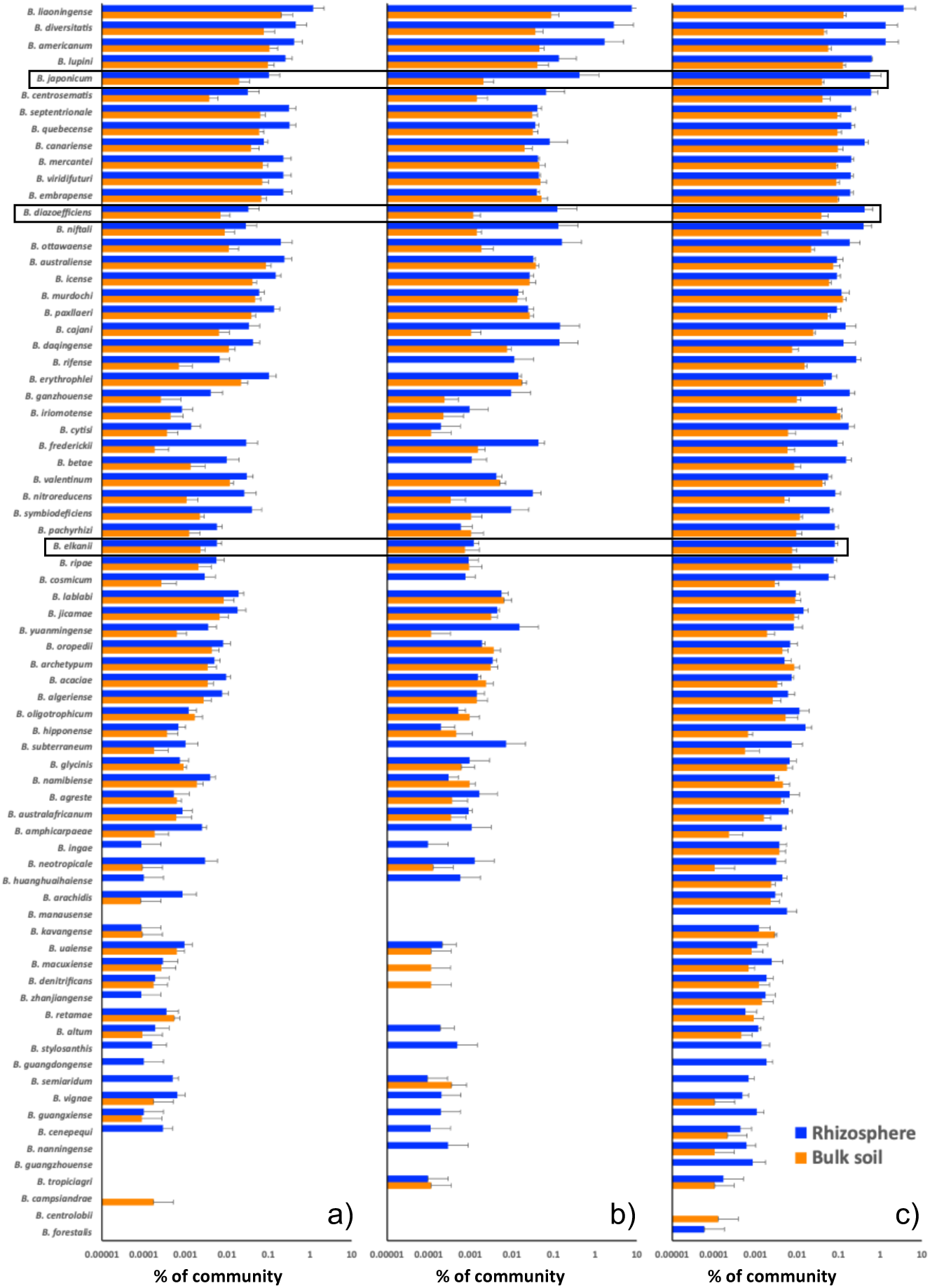
*Bradyrhizobium* species population distributions across bulk soil and rhizosphere at the three field sites. The three species obtained from nodules are indicated by boxes. The x axis reports the percentage of all reads in the sample on a log scale.

## Discussion

*Bradyrhizobium* are known to proliferate in the soybean rhizosphere. While soybean nodules mostly contain *B. japonicum, B. diazoefficiens* or *B. elkanii* (Mortuza et al 2022), over 80 species described to date have been associated with soils, and the genus is predicted to comprise over 800 species (Ormeño-Orrillo and Martinez-Romero 2019). Studies of population dynamics at the species level has not been possible without isolating and identifying large numbers of bacteria from the rhizosphere because culture independent studies have relied on short reads that do not allow allocation to species. Recent advances in ONT provide greater fidelity together with increased numbers of reads (Brown et al 2017) (Nygaard et al 2020), so we employed ONT’s sequencing platform to obtain high numbers of full 16S rRNA gene reads to reveal population dynamics of *Bradyrhizobium* and the associated bacterial communities in bulk soil, rhizosphere, and nodules from different field sites. The data obtained enabled us to determine relative quantities of 10,855 species in bulk soil and rhizosphere, including 74 *Bradyrhizobium* species.

### *Bradyrhizobium* species are successful in soils in the absence of nodulating partners

Bulk soil contained abundant *Bradyrhizobium*, in agreement with a recent meta-study of soils across the globe (Delgado-Baquerizo et al 2018). We found over 70 species in each of the three soils. It is fascinating that so many *Bradyrhizobium* species maintained populations in soils of varying pH and ionic content. While the overall bacterial community was affected by soil pH, as shown by beta diversity, the *Bradyrhizobium* composition was similar (Fig 1 c,d; 6). Soil chemical properties did not have any effect on *Bradyrhizobium* species composition or prevalence. Limitations of inorganic nutrients such as phosphorus, potassium, and of organic matter in site 6 did not affect the *Bradyrhizobium* species prevalence. This indicates that the genus is well adapted to the soil environment, and able to thrive under a range of physico-chemical conditions. The acidic site 8 contained a few more *Bradyrhizobium* species at low prevalence, suggesting that some species do exhibit niche preference. While most of the species found have been reported from nodules of diverse legumes, the field sites have not hosted any legumes other than soybeans for decades. While these *Bradyrhizobium* are able to nodulate compatible leguminous hosts, they maintained substantial populations in the soil. For example, *B. americanum* nodulates *Centrosema*, a subtropical leguminous vine that did not occur in any of the fields sampled (Ramírez-Bahena et al 2016). This points to a nodule-independent lifestyle of the genus.

### Soybean rhizosphere supports populations of diverse *Bradyrhizobium* and other nodulating fixers

*Bradyrhizobium* responded positively to the growing roots since all but a few bulk soil species were enriched in the rhizosphere. This indicates that roots provide an environment supportive of most *Bradyrhizobium* species occurring in bulk soil. This is likely due to nutrients in soybean root exudates (Sugiyama 2019). *Bradyrhizobium* was the most predominant genus in rhizosphere in all sites sampled. The predominant species were *B. liaoningense, B. americanum,* and *B. diversitatus,* but none of them were found in nodules. A diversity of other nitrogen-fixing nodulating genera including *Rhizobium* and *Mesorhizobium* were also enriched in the rhizosphere. In contrast, free-living nitrogen fixers such as *Azospirillum* and *Paenibacillus,* appeared suppressed in the soybean rhizosphere, as were many taxa indicated in Fig 2. Our findings suggest that root exudates play a key role in shaping bacterial diversity, in agreement with many previous reports (Park et al 2023), (Munoz-Ucros et al 2021). Growing soybean roots release a complex exudate into the surrounding soil. These exudates include amino acids, sugars, and organic acids (Tawaraya et al 2014), (Tantriani et al 2020), many of which are chemoattractants of *Bradyrhizobium* (Sandhu et al 2023). Roots also secrete isoflavones that induce nod factor formation in selected *Bradyrhizobium* (Sandhu et al 2021). Collectively, these data indicate that soybean roots support a wide array of associative nitrogen-fixing taxa rather than the select group able to nodulate them. Yet, entry of fixers into root hairs to populate nodules is clearly governed by factors that are species and strain-specific (Poole et al 2018).

### Nodulation is not driven by mere abundance of nodulator

The most predominant species of *Bradyrhizobium* from bulk soil and rhizosphere, *B. lianongense, B. americanum, and B. diversitatus*, did not nodulate soybean roots in these sites. While *B. lianongense* constituted 4.1% of the rhizosphere community, the nodulating *B. japonicum*, *B. elkanii,* and *B. diazoefficiens* constituted 0.36, 0.028 and 0.19 % of the rhizosphere community. While *B. americanum* was originally isolated from *Centrosema* (Ramírez-Bahena et al 2016), *B. diversitatus* from *Glycine clandestine* (Klepa et al 2021), and *B. lianongense* was isolated from a soybean plant in China (Xu et al 1995). Yet these most predominant *Bradyrhizobium* were not found in any of the nodules we sampled. The low prevalence of successful versus unsuccessful nodulators indicates that nodulation is not driven primarily by nodulator abundance. We speculate that the specificity of the three nodulators for the local soybean cultivars and soil and climate conditions was greater than *B*. *liaoningense*. This indicates that the gatekeeper functions of crosstalk between root and bacterium play a dictating role in entry of compatible *Bradyrhizobium* into the root cortical cells. Prevalence of species in nodules varied by site, with site 6 containing primarily *B. elkanii*. While *B. diazoefficiens* is known as the preferred nodulator and was present in all bulk soils and rhizosphere samples, we found it in only one nodule in the acidic site 8. The steps leading to successful entry of *Bradyrhizobium* into root cortical cells under field conditions may be affected by multiple factors, including soil chemistry, species interactions, soybean cultivar and strain-specific traits.

### Correlation of other predominant rhizosphere taxa with *Bradyrhizobium*

A variety of taxa aside from *Bradyrhizobium* became predominant in the rhizosphere. This included various associative nitrogen-fixing Hyphomicrobiales (Alphaproteobacteria) and Burkholderiales (Betaproteobacteria). It also included non-nitrogen-fixing taxa such as *Variovorax* (Burkholderiales), which comprised 9.5 % of the rhizosphere community. The soybean rhizosphere supported population success of this wider group of Alpha - and Betaproteobacteria by enriching them from bulk soil. In contrast, the bulk of classic soil bacteria such as Bacilli (Firmicutes), Actinobacteria and Acidobacteria decreased in population density. The Rhizobiaceae (*Rhizobium, Sinorhizobium, Neorhizobium, Ensifer,* and *Shinella*) and Devosiaceae (*Devosia*) correlated positively with Nitrobacteraceae (*Bradyrhizobium*), except in site 6. This positive correlation indicates that these nitrogen-fixing taxa shared this niche cooperatively, regardless of nodulation potential. Lack of correlation in site 6 indicated that physico-chemical factors, such as phosphate and potassium, appear to impact interactions among some of these taxa and the root environment. While the acidophilic Burkholderiales (Burkholderia, Paraburkholderia and Trinickia) were more prevalent in acidic site 8, they correlated positively with the Hyphomicrobiales (*Bradyrhizobium, Rhizobium, Sinorhizobium, Neorhizobium, Ensifer, Shinella,* and *Devosia*) in all three sites. This indicates cooperation among these diverse nitrogen-fixing bacteria in the rhizosphere and also in bulk soil. Whether these non-nodulating nitrogen fixers merely take advantage of the nutrient rich rhizosphere or benefit the plant or associated microbiota is unknown. One of the Burkholderiales, *Variovorax*, has been found to be an endophyte in soybean nodules (Mayhood and Mirza 2021). *V. paradoxus* and *V. gossypii*, the two predominant species found (Supplemental Table 1) are known as plant-supporting symbiont with plant growth-promoting properties (Han et al 2011). They also enhance nodulation and stress responses in *Medicago* (Flores-Duarte et al 2022). At least some of these positively correlating taxa would probably benefit the soybean plant by promoting nodulation or other growth-promoting functions. Studies of the specific roles of these diverse rhizosphere bacteria are needed to better understand rhizosphere functioning.

Our data show that field grown soybeans are surrounded by a diversity of *Bradyrhizobium* and other associative nitrogen fixers when planted in bulk soil, and that their roots support development of a rhizosphere rich in diverse associative nitrogen fixers. Only three species of one genus of this diverse rhizosphere microbiota nodulated the soybeans. The reasons underlying this broad growth supporting effect of soybean roots is unclear, as are possible positive and negative roles of these rhizosphere microbiota. Full 16S rRNA gene sequences obtained using ONT-MinION allowed allocation to species level but were insufficient to distinguish among strains. While we found evidence for soybean nodulating species B*. japonicum, B. elkanii, and B. diazoefficiens* in the bulk soil, it is unclear what proportion of these are nodulating strains. If the bulk soil B*. japonicum, B. elkanii, and B. diazoefficiens* include nodulating strains, then addition of microbial inoculants would be redundant. *Bradyrhizobium* was the most abundant genus in the bulk soil and rhizosphere in all three sites, in agreement with the findings from soils worldwide (Delgado-Baquerizo et al 2018). Our results showed that many species of the genus maintain populations in diverse cultivated soils, even in the long-term absence of potential nodulating partners.

## Acknowledgements

Sukhvir Kaur Sarao was supported by a National Science Foundation assistantship. We thank Armaan Kaur Sandhu, Muhammad Yasir Afzal, and Johnathan Orosz for their critical reading of the manuscript. This material is based upon work supported by the National Science Foundation/EPSCoR RII Track-1: Building on the 2020 Vision: Expanding Research, Education and Innovation in South Dakota, Award OIA-1849206 and by the South Dakota Board of Regents. This material is based upon work conducted using the South Dakota State University Functional Genomics Core Facility (RRID:SCR_023786). The Genomics Sequencing Facility (RRID:SCR_023959) is partially funded by the South Dakota Agricultural Experimental Station and the National Institute of General Medical Sciences of the National Institutes of Health under Award Number P20GM135008. The content is solely the responsibility of the authors and does not necessarily represent the official views of the National Institutes of Health.

## References

Altschul SF, Gish W, Miller W, Myers EW, Lipman DJ (1990). Basic local alignment search tool. Journal of molecular biology 215: 403–410.

Brown BL, Watson M, Minot SS, Rivera MC, Franklin RB (2017). MinION™ nanopore sequencing of environmental metagenomes: a synthetic approach. Gigascience 6: gix007.

Buhian WP, Bensmihen S (2018). Mini-review: nod factor regulation of phytohormone signaling and homeostasis during rhizobia-legume symbiosis. Frontiers in plant science 9: 1247.

Delgado-Baquerizo M, Oliverio AM, Brewer TE, Benavent-González A, Eldridge DJ, Bardgett RD et al (2018). A global atlas of the dominant bacteria found in soil. Science 359: 320–325.

Du X, Liu N, Yan B, Li Y, Liu M, Huang Y (2024). Proximity-based defensive mutualism between *Streptomyces* and *Mesorhizobium* by sharing and sequestering iron. The ISME Journal 18: wrad041.

Flores-Duarte NJ, Pérez-Pérez J, Navarro-Torre S, Mateos-Naranjo E, Redondo-Gómez S, Pajuelo E et al (2022). Improved *Medicago sativa* nodulation under stress assisted by *Variovorax* sp. endophytes. Plants 11: 1091.

Galet J, Deveau A, Hôtel L, Frey-Klett P, Leblond P, Aigle B (2015). *Pseudomonas fluorescens* pirates both ferrioxamine and ferricoelichelin siderophores from *Streptomyces ambofaciens*. Applied and Environmental Microbiology 81: 3132–3141.

Ha A, Saleh-Ranstin N (2002). Symbiotical characteristics of indigenous *Sinorhizobium meliloti* strains from some Iranian soils and their variations in the different levels of salinity.

Han J-I, Choi H-K, Lee S-W, Orwin PM, Kim J, LaRoe SL et al (2011). Complete genome sequence of the metabolically versatile plant growth-promoting endophyte *Variovorax paradoxus* S110. Journal of bacteriology 193: 1183–1190.

Hassan MK, McInroy JA, Kloepper JW (2019). The interactions of rhizodeposits with plant growth-promoting rhizobacteria in the rhizosphere: a review. Agriculture 9: 142.

Herschend J, Ernst M, Koren K, Melnik AV, da Silva RR, Røder HL et al (2022). Metabolic Profiling of Interspecies Interactions During Sessile Bacterial Cultivation Reveals Growth and Sporulation Induction in *Paenibacillus amylolyticus* in Response to *Xanthomonas retroflexus*. Frontiers in cellular and infection microbiology 12: 805473.

Howard JB, Rees DC (1996). Structural basis of biological nitrogen fixation. Chemical reviews 96: 2965–2982.

Klepa MS, Ferraz Helene LC, O’Hara G, Hungria M (2021). *Bradyrhizobium agreste* sp. nov., *Bradyrhizobium glycinis* sp. nov. and *Bradyrhizobium diversitatis* sp. nov., isolated from a biodiversity hotspot of the genus *Glycine* in Western Australia. International Journal of Systematic and Evolutionary Microbiology 71: 004742.

Koirala A, Brözel VS (2021). Phylogeny of nitrogenase structural and assembly components reveals new insights into the origin and distribution of nitrogen fixation across bacteria and archaea. Microorganisms 9: 1662.

Li QQ, Wang ET, Chang YL, Zhang YZ, Zhang YM, Sui XH et al (2011). *Ensifer sojae* sp. nov., isolated from root nodules of *Glycine max* grown in saline-alkaline soils. International journal of systematic and evolutionary microbiology 61: 1981–1988.

Lu Y, Zhou G, Ewald J, Pang Z, Shiri T, Xia J (2023). MicrobiomeAnalyst 2.0: comprehensive statistical, functional and integrative analysis of microbiome data. Nucleic Acids Research: gkad407.

Lyng M, Jørgensen JP, Schostag MD, Jarmusch SA, Aguilar DK, Lozano-Andrade CN et al (2024). Competition for iron shapes metabolic antagonism between B*acillus subtilis* and *Pseudomonas marginalis*. The ISME journal 18: wrad001.

Mayhood P, Mirza BS (2021). Soybean root nodule and rhizosphere microbiome: Distribution of rhizobial and nonrhizobial endophytes. Applied and Environmental Microbiology 87: e02884–02820.

Mortuza MF, Djedidi S, Ito T, Agake S-i, Sekimoto H, Yokoyama T et al (2022). Genetic and Physiological Characterization of Soybean-Nodule-Derived Isolates from Bangladeshi Soils Revealed Diverse Array of Bacteria with Potential Bradyrhizobia for Biofertilizers. Microorganisms 10: 2282.

Munoz-Ucros J, Zwetsloot MJ, Cuellar-Gempeler C, Bauerle TL (2021). Spatiotemporal patterns of rhizosphere microbiome assembly: From ecological theory to agricultural application. Journal of Applied Ecology 58: 894–904.

Mylona P, Pawlowski K, Bisseling T (1995). Symbiotic nitrogen fixation. The Plant Cell 7: 869.

Nakei MD, Venkataramana PB, Ndakidemi PA (2022). Soybean-nodulating rhizobia: Ecology, characterization, diversity, and growth promoting functions. Frontiers in Sustainable Food Systems 6: 824444.

Nygaard AB, Tunsjø HS, Meisal R, Charnock C (2020). A preliminary study on the potential of Nanopore MinION and Illumina MiSeq 16S rRNA gene sequencing to characterize building-dust microbiomes. Scientific Reports 10: 3209.

ONT ONT (2023). EPI2ME Desktop Software.

Ormeño-Orrillo E, Martinez-Romero E (2019). A genomotaxonomy view of the *Bradyrhizobium* genus. Frontiers in microbiology 10: 1334.

Park I, Seo Y-S, Mannaa M (2023). Recruitment of the rhizo-microbiome army: assembly determinants and engineering of the rhizosphere microbiome as a key to unlocking plant potential. Frontiers in Microbiology 14: 1163832.

Parker MA (2015). The spread of *Bradyrhizobium* lineages across host legume clades: from *Abarema* to *Zygia*. Microbial ecology 69: 630–640.

Poole P, Ramachandran V, Terpolilli J (2018). Rhizobia: from saprophytes to endosymbionts. Nature Reviews Microbiology 16: 291–303.

Ramírez-Bahena MH, Flores-Félix JD, Chahboune R, Toro M, Velázquez E, Peix A (2016). *Bradyrhizobium centrosemae* (symbiovar centrosemae) sp. nov., B*radyrhizobium americanum* (symbiovar phaseolarum) sp. nov. and a new symbiovar (tropici) of *Bradyrhizobium viridifuturi* establish symbiosis with *Centrosema* species native to America. Systematic and applied microbiology 39: 378–383.

Sandhu AK, Subramanian S, Brözel VS (2021). Surface properties and adherence of *Bradyrhizobium diazoefficiens* to *Glycine max* roots are altered when grown in soil extracted nutrients. Nitrogen 2: 461–473.

Sandhu AK, Brown MR, Subramanian S, Brözel VS (2023). *Bradyrhizobium diazoefficiens* USDA 110 displays plasticity in the attachment phenotype when grown in different soybean root exudate compounds. Frontiers in Microbiology 14: 1190396.

Schoch CL, Ciufo S, Domrachev M, Hotton CL, Kannan S, Khovanskaya R et al (2020). NCBI Taxonomy: a comprehensive update on curation, resources and tools. Database (Oxford) 2020.

Schultze M, Kondorosi A (1998). Regulation of symbiotic root nodule development. Annual review of genetics 32: 33–57.

Stougaard J (2000). Regulators and regulation of legume root nodule development. Plant physiology 124: 531–540.

Subramanian S, Stacey G, Yu O (2006). Endogenous isoflavones are essential for the establishment of symbiosis between soybean and *Bradyrhizobium japonicum*. The Plant Journal 48: 261–273.

Sugiyama A (2019). The soybean rhizosphere: Metabolites, microbes, and beyond—A review. Journal of advanced research 19: 67–73.

Sun H, Jiang S, Jiang C, Wu C, Gao M, Wang Q (2021). A review of root exudates and rhizosphere microbiome for crop production. Environmental Science and Pollution Research 28: 54497–54510.

Tamura K, Stecher G, Kumar S (2021). MEGA11: molecular evolutionary genetics analysis version 11. Molecular biology and evolution 38: 3022–3027.

Tantriani, Shinano T, Cheng W, Saito K, Oikawa A, Purwanto BH et al (2020). Metabolomic analysis of night-released soybean root exudates under high-and low-K conditions. Plant and Soil 456: 259–276.

Tawaraya K, Horie R, Shinano T, Wagatsuma T, Saito K, Oikawa A (2014). Metabolite profiling of soybean root exudates under phosphorus deficiency. Soil science and plant nutrition 60: 679–694.

Team R (2021a). RStudio: Integrated Development for R.

Team TRC (2021b). R: A language and environment for statistical computing. R Foundation for Statistical Computing.

Tong Z, Sadowsky MJ (1994). A selective medium for the isolation and quantification of *Bradyrhizobium japonicum* and *Bradyrhizobium elkanii* strains from soils and inoculants. Applied and environmental microbiology 60: 581–586.

White LJ, Brözel VS, Subramanian S (2015). Isolation of rhizosphere bacterial communities from soil. Bio-protocol 5: e1569–e1569.

Wood DE, Lu J, Langmead B (2019). Improved metagenomic analysis with Kraken 2. Genome Biology 20: 257.

Xu L, Ge C, Cui Z, Li J, Fan H (1995). *Bradyrhizobium liaoningense* sp. nov., isolated from the root nodules of soybeans. International Journal of Systematic and Evolutionary Microbiology 45: 706–711.

