## Supplemental Figure 1 for "*Bradyrhizobium* and the soybean rhizosphere: Species level bacterial population dynamics in established soybean fields, rhizosphere and nodules"

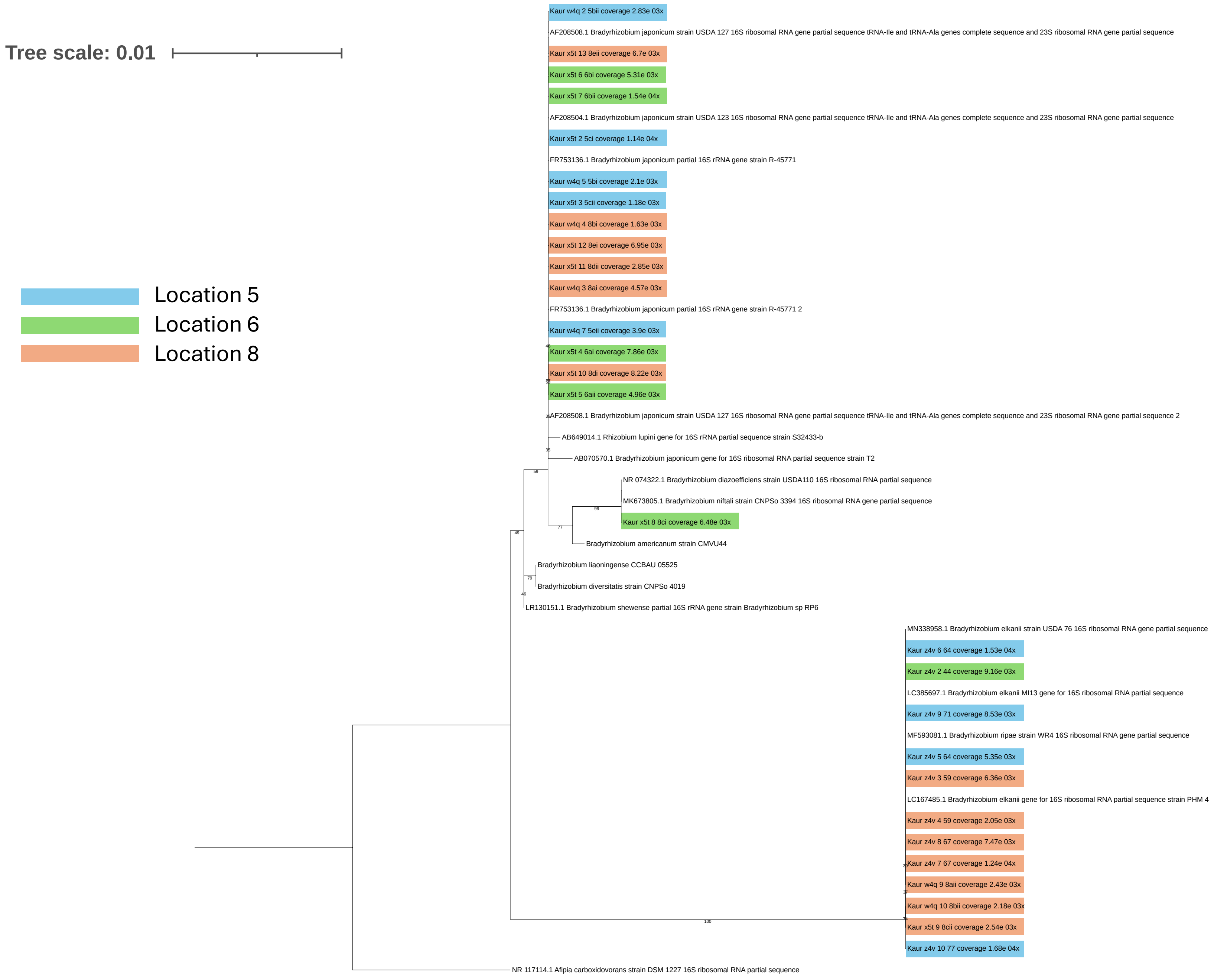


Figure S1: Maximum likelihood dendrogram of *Bradyrhizobium* nodule isolates and reference sequences obtained from sequenced genomes, using full 16SrRNA gene sequences.
